# nanoPhos enables ultra-sensitive and cell-type resolved spatial phosphoproteomics

**DOI:** 10.1101/2025.05.29.656770

**Authors:** Denys Oliinyk, Tim Heymann, Lukas Henneberg, Anastasiya Bardziukova, Marvin Thielert, Jeppe Kjærgaard, Maximilian Zwiebel, Sabine Vasconez, Marc Oeller, Andreas Metousis, Enes Ugur, Edwin H. Rodriguez, Florian A. Rosenberger, Simon Schallenberg, Frederick Klauschen, Matthias Mann

**Affiliations:** Department of Proteomics and Signal Transduction, Max Planck Institute of Biochemistry, Martinsried, Germany; NNF Center for Basic Metabolic Research, University of Copenhagen, Copenhagen, Denmark; Department of Medical Biochemistry and Biophysics, Karolinska Institutet, Solna, Sweden; Institute of Pathology, Charité Universitätsmedizin Berlin, Berlin, Germany; Institute of Pathology, LMU Munich, Munich, Germany

**Author notes:** These authors contributed equally (Denys Oliinyk, Tim Heymann).

## Abstract

Mass spectrometry (MS)-based phosphoproteomics has transformed our understanding of cell signaling, yet current workflows face limitations in sensitivity and spatial resolution at sub-microgram inputs. Here, we present nanoPhos, a robust method that extends phosphoproteomics to nanogram scale, making it compatible with cell-type-resolved spatial analysis. It employs loss-less solid phase extraction capture (SPEC) for sample preparation, followed by automated phosphopeptide enrichment using Fe(III)-NTA cartridges. nanoPhos identifies over 57,000 unique phosphorylation sites from 1 µg cell lysate and over 4,000 from only 10 ng, a hundred-fold improvement from recent protocols. Combined with Deep Visual Proteomics (DVP), it enables region- and cell-type resolved phosphoproteomics of mouse brain tissue with spatial fidelity and a depth of 13,000 phosphosites from only 1000 cell shapes. This establishes nanoPhos as a versatile and ultra-sensitive platform that extends DVP to post-translational modifications and opens up for cell-type-specific signaling analysis in intact tissue.

## Introduction

MS-based phosphoproteomics has become a powerful tool for mapping signaling networks at proteome scale^1^. Over the past decade, advances in MS instrumentation, sample preparation, and data analysis have enabled increasingly deep and quantitative analysis of protein phosphorylation from milligrams of input material to the microgram range^2–4^. Streamlined workflows such as EasyPhos minimized manual sample processing and enabled high-throughput phosphoproteome profiling, facilitating large-scale biological applications^5–9^. Recently, the µPhos and other protocols extended these capabilities to low-microgram inputs through optimized phosphopeptide enrichment in 96-well formats^10–12^. Together such advances enabled the in-depth mapping of phosphorylation networks in cells and tissues and their functional characterization in multi-condition perturbation studies and drug profiling^13^

However, several challenges have prevented further miniaturization of phosphoproteomics workflows towards nanogram-scale input amounts. These include reliance on relatively large processing volumes to maintain optimal peptide concentrations for bead-based in-solution workflows and elaborate conditions for the optimal enrichment of phosphorylated peptides. Furthermore, detergents, while important for cell lysis, compromise enrichment efficiency^12^. Addressing these challenges could achieve the sensitivity needed to extend phosphoproteomics to rare cell populations isolated by fluorescence activated cell sorting (FACS), archived formalin-fixed paraffin-embedded (FFPE) clinical specimens, microdissected tissue regions, and ultimately to cell-type-resolved spatial contexts.

We recently developed Deep Visual Proteomics (DVP), combining high-content imaging, AI-driven cell classification, and laser microdissection with ultra-sensitive mass spectrometry to profile proteomes at single-cell type resolution^14^. So far, DVP has been limited to measurements of protein abundance only. Extending it to cell signaling would open a new biological dimension - how signaling networks operate within their native spatial and cellular context^15–18^. Here we describe nanoPhos, a phosphoproteomics workflow that addresses these challenges, enabling nanogram inputs. We apply nanoPhos to cultured cells, primary tissue, and laser-microdissected cell populations, demonstrating deep, cell-type-resolved phosphoproteomics that captures signaling *in situ*.

## Results

### A phosphoproteomics workflow for nanogram-scale samples

The core principle of nanoPhos is minimizing sample volumes at every step, from lysis through enrichment. Conventional phosphoproteomics workflows require microliter-scale volumes for bead-based enrichment, causing proportionally greater adsorptive and transfer losses at low inputs. We integrated Solid-Phase Extraction and Capture (SPEC), which concentrates proteins into nanoliter-scale volumes within a standard pipette tip, enabling efficient digestion with fast kinetics and broad detergent compatibility (**Fig. 1**). The SPEC workflow is described in detail in a companion paper^19^. This allows strong lysis conditions without the sample losses typically associated with detergent removal. We routinely use 2% SDC in nanoPhos, ensuring that phosphatase activity is quenched and that even membrane and cytoskeletal proteins are fully solubilized.

**Figure 1.**
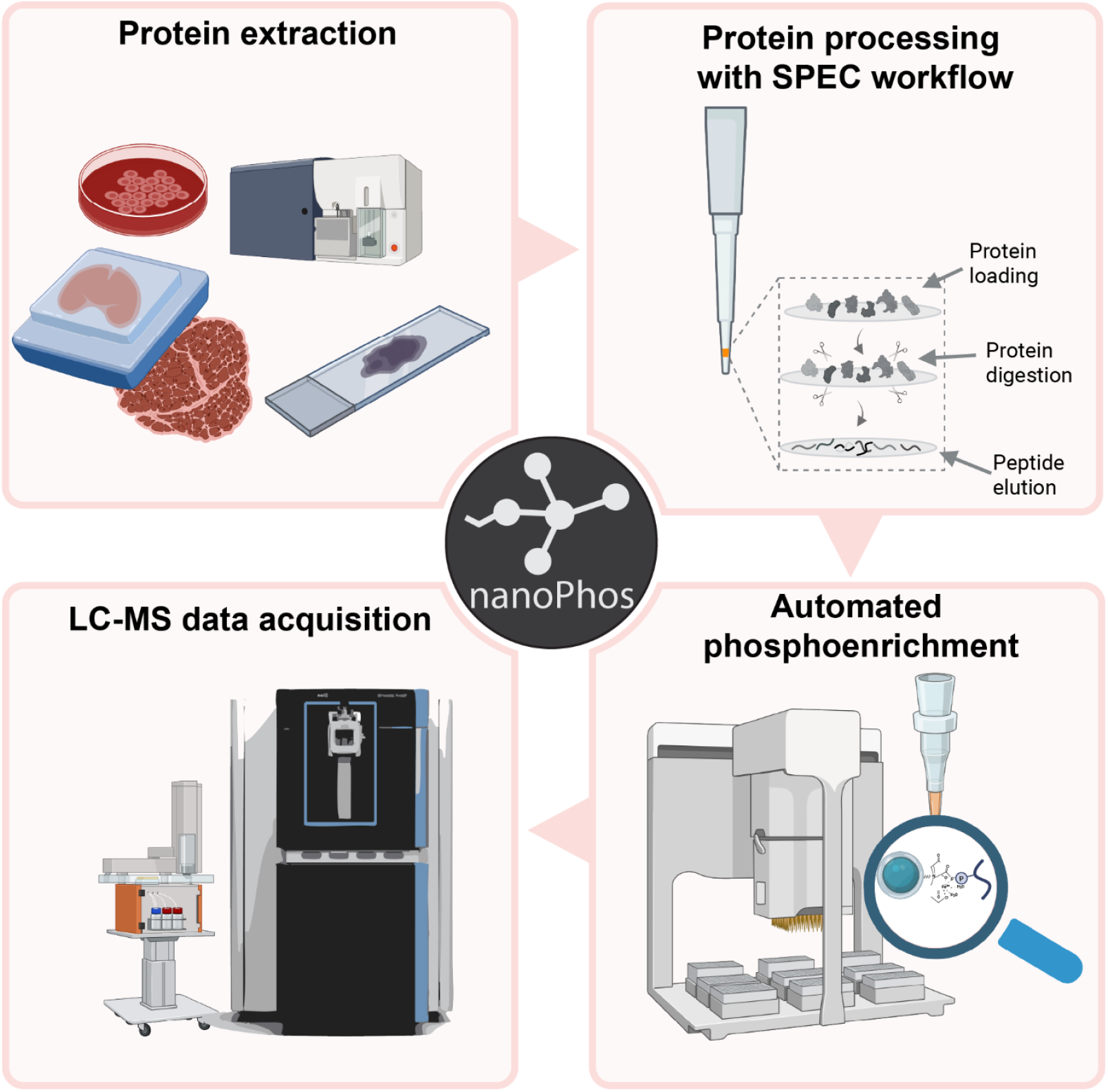
Design of the nanoPhos workflow. The protocol enables efficient detergent-based lysis, proteolytic digestion, and peptide cleanup in nanoliter volumes. Zero dead-volume phosphopeptide enrichment on Fe(III)-NTA cartridges eliminates absorptive loss and allows for direct injection of enriched phosphopeptides into LC-MS/MS.

Peptides are eluted in low volume and directly subjected to zero-dead-volume phosphopeptide enrichment on a robotic platform (AssayMAP Bravo), using Fe(III)-NTA cartridges; this limits the phosphopeptide enrichment volumes to the effective volume of the cartridge and eliminates the transfer losses that limit conventional bead-based protocols. Each module - lysis, digestion, enrichment - can be independently optimized and suited to the users’ needs, thus maintaining full compatibility with high-throughput formats and ultra-low input amounts (**Methods**). For instance, addition of 200 mM NaCl to the enrichment buffer, made possible by the SPEC workflow, substantially reduces co-enrichment of non-phosphorylated peptides, achieving selectivity exceeding 60% even at the lowest inputs (**Suppl. Fig. 1a**).

Enriched phosphopeptides are eluted from Fe(III)-NTA cartridges directly into Evotips for seamless integration with downstream data-independent acquisition (DIA)-MS on an Orbitrap – Astral platform, avoiding most of the transfer steps and associated sample losses. The high scan speed of the Astral analyzer enables narrow-window or variable window DIA schemes that maximize precursor selectivity and quantitative accuracy from limited phosphopeptide yields^20,21^. The entire process, from cell or tissue lysate to MS-ready sample, takes about two hours and supports diverse input types, as shown below.

### Benchmarking nanoPhos sensitivity and quantitative performance

To evaluate nanoPhos, we performed phosphopeptide enrichment from HeLa cell lysates in both EGF-stimulated and untreated conditions (**Fig. 2a**). Starting with 1 µg of input - at the low edge of literature reports - nanoPhos identified more than 57,000 unique phosphorylation sites, on 4,868 proteins, representing one of the deepest single-run phosphoproteomes reported to date (**Fig. 2b**). This depth covered more than 60% of the sites likely to be functional (‘functional score’ > 0.5) in the Ochoa et al. functional phosphoproteome^22^ (**Suppl. Fig. 1b**). Coverage remained at 36,799 sites for 200 ng of input and still more than 4,000 sites on 961 proteins at only 10 ng of which about 2,000 were Class I sites (localization probability > 75%).

**Figure 2.**
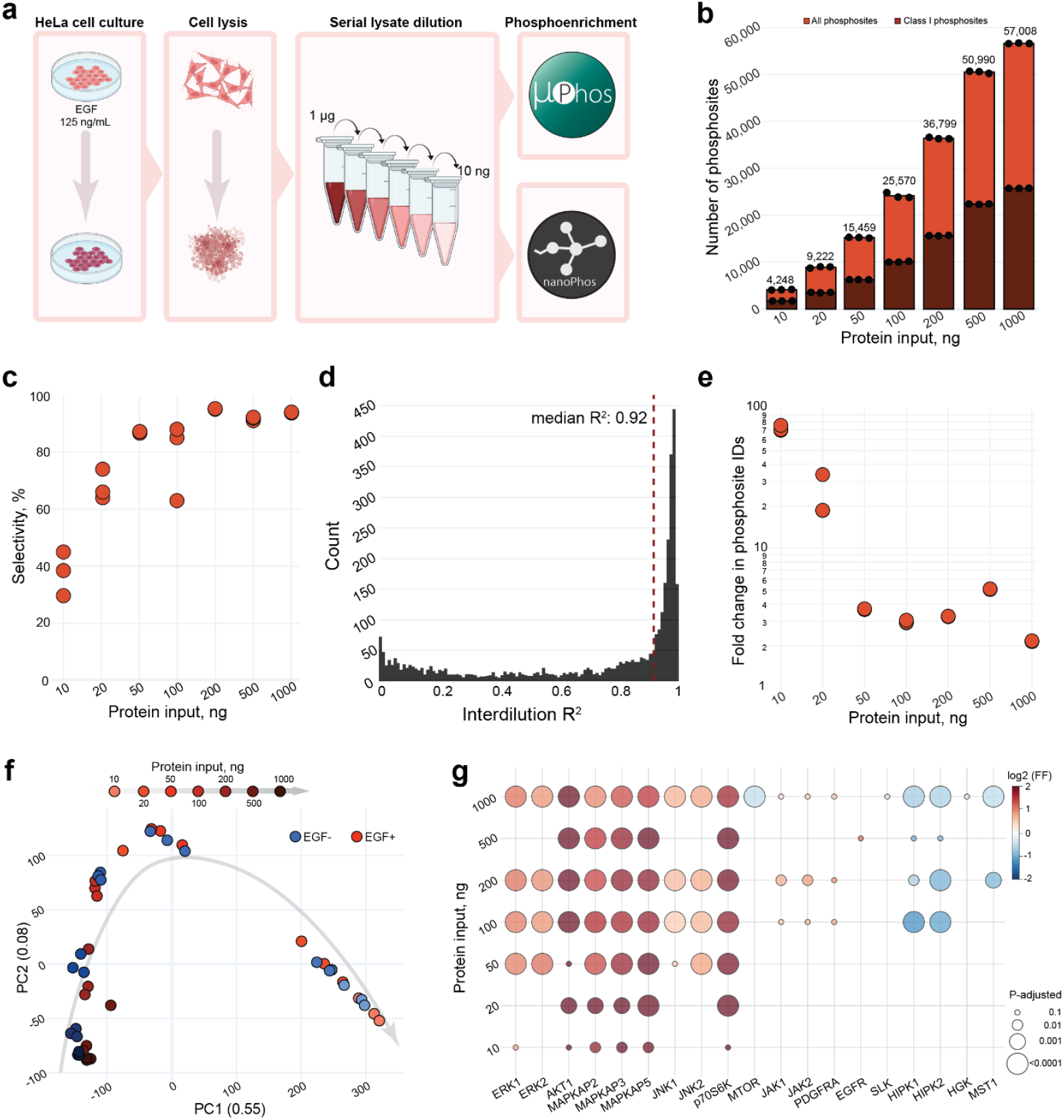
nanoPhos achieves orders of magnitude improvement in sensitivity. **a** Schematic of cell line- based phosphoproteomics workflow. **b** Number of identified phosphosites and Class I phosphosites (localization probability > 0.75, darker color) as a function of HeLa protein input (n = 3 per condition). **C** Selectivity of phosphopeptide enrichment across the dilution series (n = 3 per condition). **d** Coefficient of determination (R^2^) for the intensity of every phosphosite identified in > 3 dilution points as a function of input amount. **e** Fold change difference in number of phosphosite identifications between nanoPhos and µPhos. **f** PCA of EGF-treated (EGF+) and untreated (EGF-) HeLa samples across protein input amounts (n = 3 per condition). **g** Kinase substrate enrichment a for t-test significant phosphosites (Adj. P-value < 0.05, |log2FC| > 0.585) between EGF-treated and untreated samples (n = 3 per condition). Bubble size represents adjusted p-value of kinase enrichment analysis (Adj. P-value < 0.05) and color indicates log-transformed enrichment score (positive for enriched kinases, negative for depleted).

To assess quantitative accuracy, we asked how faithfully phosphosite intensities scale with input amount. For each phosphorylation site detected in at least three dilution points, we fitted a linear regression and calculated inter-dilution coefficients of determination (R^2^). A median R^2^ of 92% demonstrates high quantitative precision (**Fig. 2d**). Interreplicate coefficients of variation were ∼18% (**Suppl. Fig. 1c**) with no significant changes between dilutions. Furthermore, phosphopeptide selectivity remained above 75% for protein inputs above 10ng (**Fig. 2c**), confirming that sensitivity gains did not compromise enrichment quality.

We directly compared nanoPhos with the recently published µPhos platform, using the Astral mass spectrometer for both protocols. The differences were most pronounced at 10 ng, where nanoPhos identified nearly a hundred times more phosphosites. Between 100 ng and 1 µg, it delivered a consistent four-fold increase (**Fig. 2e**). nanoPhos-enriched peptides showed higher GRAVY indices, indicating improved detection of hydrophobic sequences that are often lost in conventional workflows due to adsorptive losses (**Suppl. Fig. 1d**).

EGF signaling in HeLa cells has been used for over 20 years as a prototypical signaling system for evaluating phosphoproteomics technologies^21,23,24^. This system is well-characterized and remains a demanding test of quantitative fidelity, dynamic range, and biological interpretability, especially at low input levels. We treated HeLa cells with EGF for 15 min and performed a dilution series from 1 µg to 10 ng. Principal component analysis (PCA) clearly separated EGF-treated from untreated samples at all input levels (**Fig. 2f, Suppl. Fig. 1e-k**). The number of identified phosphosites scaled proportionally with input, ranging from 3,412 sites at 10 ng to 65,718 sites at 1 µg (**Suppl. Fig. 1l**). Kinase enrichment analysis revealed that MAPK cascade kinases responded most strongly, including ERK1/2, p70S6K and JNK1/2 kinases (**Fig. 2g**). Receptor tyrosine kinases such as (EGFR, JAK1/2, and PDGFRA) were enriched only at higher input, consistent with their low stoichiometry.

### nanoPhos enables deep phosphoproteomics from individually-sorted cells

To extend nanoPhos to individually selected cells, we used FACS to make a dilution series of HeLa cells ranging from 3,000 to 100 cells. nanoPhos identified more than 52,000 unique phosphosites from 3,000 cells, and almost 10,000 from just 100 cells (**Fig. 3a**). Quantitative precision and selectivity remained high **(Fig. 3b, Suppl. Fig. 2a–b**). Direct comparison to µPhos showed a ten-fold advantage at 100 cells, narrowing to 3-fold at 3,000 cells (**Fig. 3c, Suppl. Fig. 2c**).

**Figure 3.**
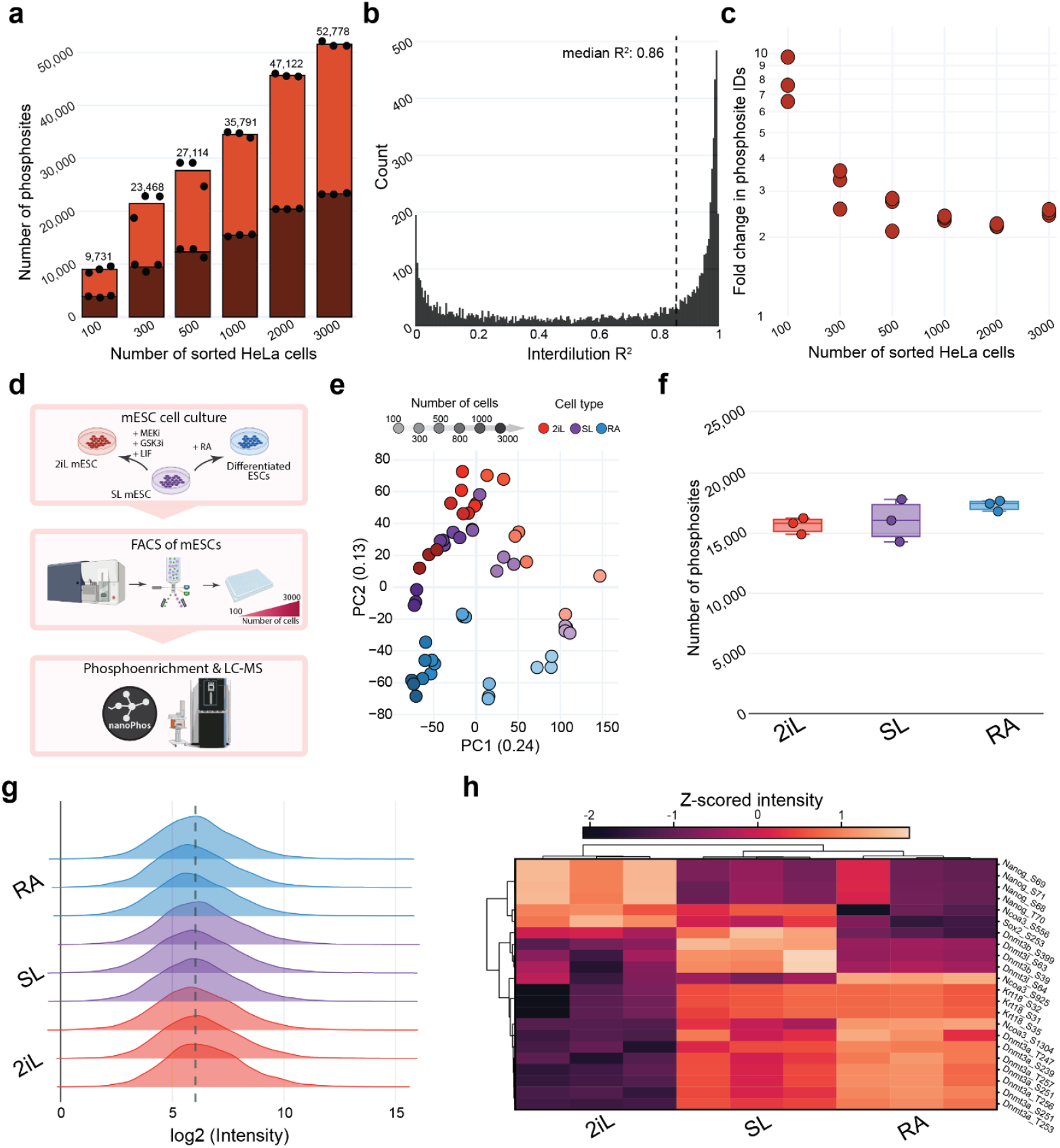
nanoPhos workflow for FACS-sorted cells and mouse embryonic stems cells. **a** Number of identified phosphosites and Class I phosphosites (localization probability > 0.75, darker color) as a function of FACS-sorted HeLa cell number (n = 3 per condition). **b** Quantitative accuracy (R^2^) across the sorted cell dilution series **c** Fold change difference in phosphosite identifications for nanoPhos versus uPhos as a function of cell number. **d** Experimental workflow for mouse embryonic stem cell (mESC) phosphoproteomics. **e** PCA of phosphoproteome data from the three mESC conditions across sorted cell numbers (n = 3 per condition). **f** Number of identified phosphosites across the three conditions of 500 sorted mESC cells (n = 3 per cell type). **g** Distribution of phosphosite intensities across the three conditions of 500 sorted mESC cells. **h** Hierarchical clustering heatmap of z-scored phosphosite intensities for significantly regulated phosphorylation sites (two-way ANOVA, FDR < 0.05) on differentiation-specific transcription factors.

To demonstrate biological applicability, we cultured mouse embryonic stem cells (mESCs) under three conditions: serum-LIF (SL) for baseline pluripotency, 2iL medium (MEK and GSK3 inhibitors plus LIF) for ground-state pluripotency, and retinoic acid (RA) for neural lineage specification (**Fig. 3d**). Cells from each condition were then FACS-sorted and processed with nanoPhos. PCA clearly separated three conditions, with PC1 reflecting cell number while PC2 distinguished between cell states (**Fig. 3e**).

Overall, we identified between 3,500 phosphosites at 100 cells and 50,000 at 3,000 cells input (**Suppl. Fig. 2d**). At 500 cells, nanoPhos identified over 15,000 phosphosites in each cell type with highly reproducible intensity distributions (**Fig. 3f-g**). Unsupervised hierarchical clustering of significantly regulated phosphosites revealed condition-specific signatures consistent with the differentiation states (**Fig. 3h**). Pluripotency factors (NANOG, SOX2, ESRRB) expectedly showed elevated phosphorylation in 2iL condition, which is required for ESC self-renewal^25^. Neural lineage markers (i.e NCOA3 S1304) emerged as upregulated in RA-treated cells, while phosphorylation events on a disordered DNMT3A region (DNMT3A S251, T253, T256, and T257) were upregulated in the SL condition, which is correlated with a nuclear localization of this protein and elevated DNA methylation levels in the SL condition^26^. This demonstrates the ability of nanoPhos to capture biologically meaningful phosphorylation differences from as few as 500 sorted cells.

### nanoPhos enables deep phosphoproteomics from tissues

We next applied nanoPhos to tissue samples, starting with mouse brain prepared as both fresh-frozen and FFPE. The latter is of particular interest given the large number of samples in biorepositories, but poses challenges due to deparaffinization and protein cross-linking. This has typically confined phosphoproteome analysis to the hundreds of microgram range so far ^27–29^.

We prepared both sample types from the same mouse brain (one hemisphere was prepared as fresh-frozen, the other was fixed with formalin and embedded in paraffin), and performed a dilution series from 10 ng to 1 µg of starting protein material. At 1 µg, nanoPhos identified 32,000 phosphosites from fresh-frozen and 25,000 from FFPE tissue. Even at 50 ng - equivalent to 50 motor neurons^30^ - we obtained over 12,000 phosphosites from fresh-frozen and 4,000 from FFPE (**Fig. 4a, Suppl. Fig. 3a-3b**). Across the full dilution series we obtained a high degree of reproducibility with median coefficient of variation at 17% - 20%. (**Fig. 4b**). At lower points of dilution series fresh-frozen tissues yielded 3-4 times more phosphosites in comparison to FFPE while at higher this difference decreased to 1.2-fold (**Suppl. Fig. 3c**).

**Figure 4.**
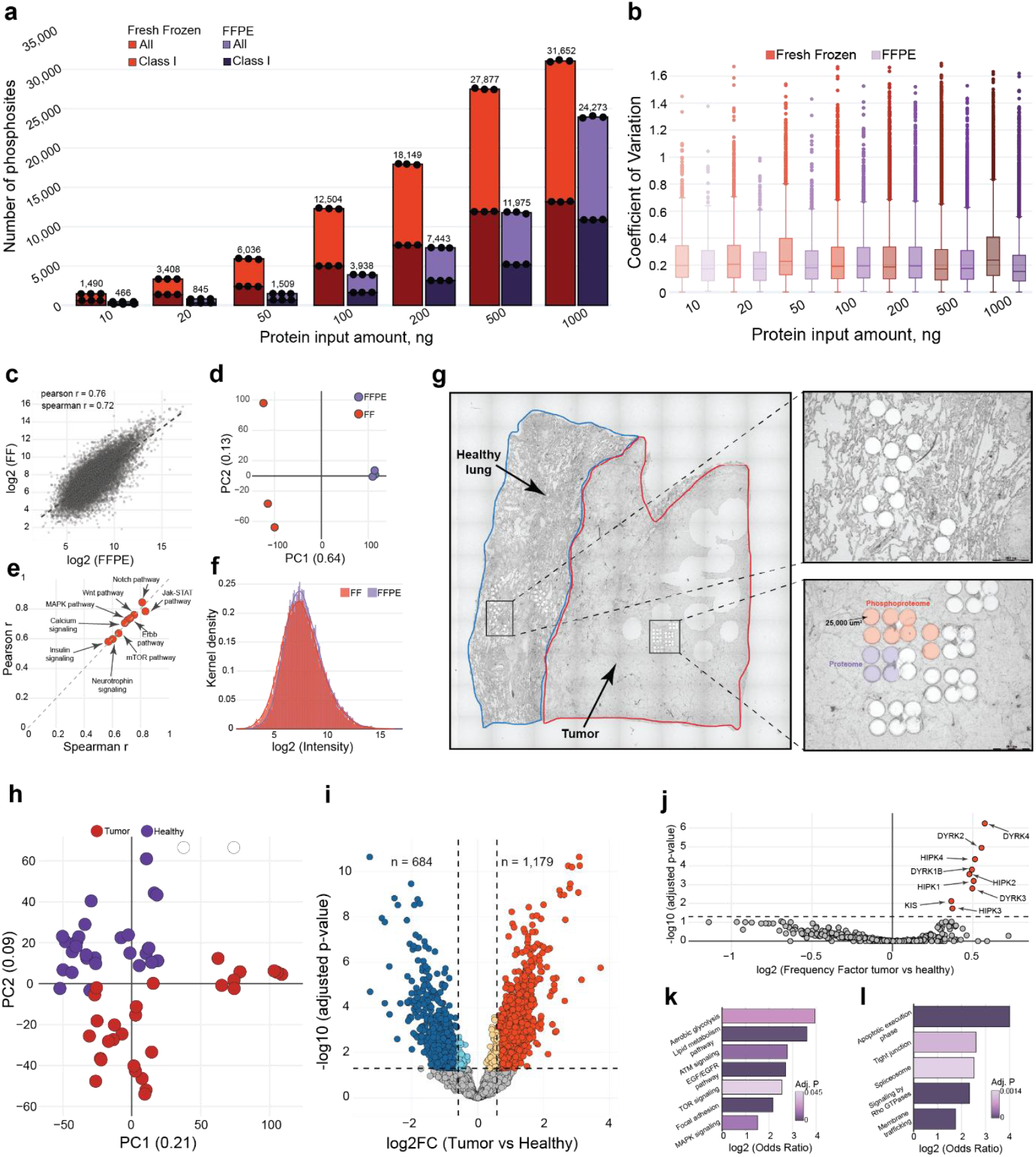
nanoPhos enables deep phosphoproteomics from fresh-frozen and FFPE tissues. **a** Number of unique and Class I phosphosites identified from fresh-frozen and FFPE mouse brain lysates as a function of protein input (n = 3). **b** Precision of label-free phosphopeptide quantification shown as coefficient of variation across workflow replicates (n = 3 per condition). **c** Correlation between fresh-frozen and FFPE phosphoproteomes at 1 µg; scatter plot of log2-transformed intensities. **d** PCA of fresh-frozen and FFPE samples for 1 µg input amount. **e** Scatter plot illustrating Pearson and Spearman correlation coefficients of major signaling pathways between fresh-frozen and FFPE tissue **f** Kernel density distribution of phosphopeptides identified in fresh-frozen and FFPE tissues. **g** Histological images of human lung tissue and adenocarcinoma sections. **h** PCA of FFPE phosphoproteome samples from adenocarcinoma (n = 28) and healthy (n = 28) regions. **i** Volcano plot (Adj. P-value < 0.05, |log2FC| > 0.585) showing differential phosphosite regulation between adenocarcinoma and healthy tissue. **j** Kinase substrate enrichment analysis of significantly regulated phosphosites. **k** GO enrichment analysis of upregulated phosphorylated proteins. Representative significantly enriched terms are shown (Adj. P-value < 0.05). **l** Same as **k** but for downregulated phosphorylated proteins.

Fresh-frozen and FFPE phosphoproteomes showed good overall correlation (Pearson *r* = 0.76 and Spearman *ρ* = 0.72) (**Fig. 4c**) while PCA revealed clear separation by sample type, reflecting systematic differences introduced by fixation and embedding (**Fig. 4d**). Importantly, major signaling pathways showed similar representation in both sample types, with high correlation scores for pathway-level phosphorylation patterns (**Fig. 4e**) with overall comparable intensity distributions (**Fig. 4f**). This indicates that while FFPE processing reduces the absolute number of retrievable phosphopeptides, it preserves both quantitative reproducibility and pathway-level representation.

Given the compatibility of nanoPhos with FFPE material, we set out to explore its applicability to spatially resolved phosphoproteomics analysis of archived clinical tissue sections. Thus, we applied nanoPhos to archived FFPE tissue from lung adenocarcinoma patients, using laser capture microdissection to isolate tumor and healthy appearing tissue regions (200,000 µm^2^ for phospho- and 100,000 µm^2^ for proteome analysis, corresponding to 500 to 2,000 cells, depending on the cell type **Fig. 4g**). This “microbulk” approach enables spatially resolved phosphoproteomics from pathologically defined areas. From these we achieved a coverage of >20,000 phosphosites and >8,000 protein groups (**Suppl. Fig. 3d**). PCA separated tumor from healthy tissue areas on phosphoproteome level and differential analysis revealed 684 significantly upregulated and 1,179 downregulated phosphosites (**Fig. 4h-i**). Kinase substrate enrichment analysis highlighted DYRK and HIPK family kinases in tumor tissue (**Fig. 4j**), consistent with their roles in cancer cell proliferation. Upregulated phosphorylated proteins were associated with aerobic glycolysis, ATM signaling, MAPK and TOR pathways (**Fig. 4k**), consistent with the metabolic and signaling alterations typically observed in *KRAS*-mutant tumors. Downregulated proteins were mapped to tight junctions, apoptotic execution phase and membrane trafficking (**Fig. 4l**).

### nanoPhos extends Deep Visual Proteomics to phosphorylation

Finally, we applied nanoPhos to cell type-resolved tissue biology by integrating it with Deep Visual Proteomics (DVP) (**Fig. 5a**). Using high-content imaging, we identified excitatory and inhibitory neurons within cortical and subcortical regions of mouse brain through multiplexed RNA-based fluorescence labelling. Cell bodies were segmented, filtered by anatomical location, and individually laser microdissected for nanoPhos analysis. We term this integrated workflow “phosphoDVP.”

**Figure 5.**
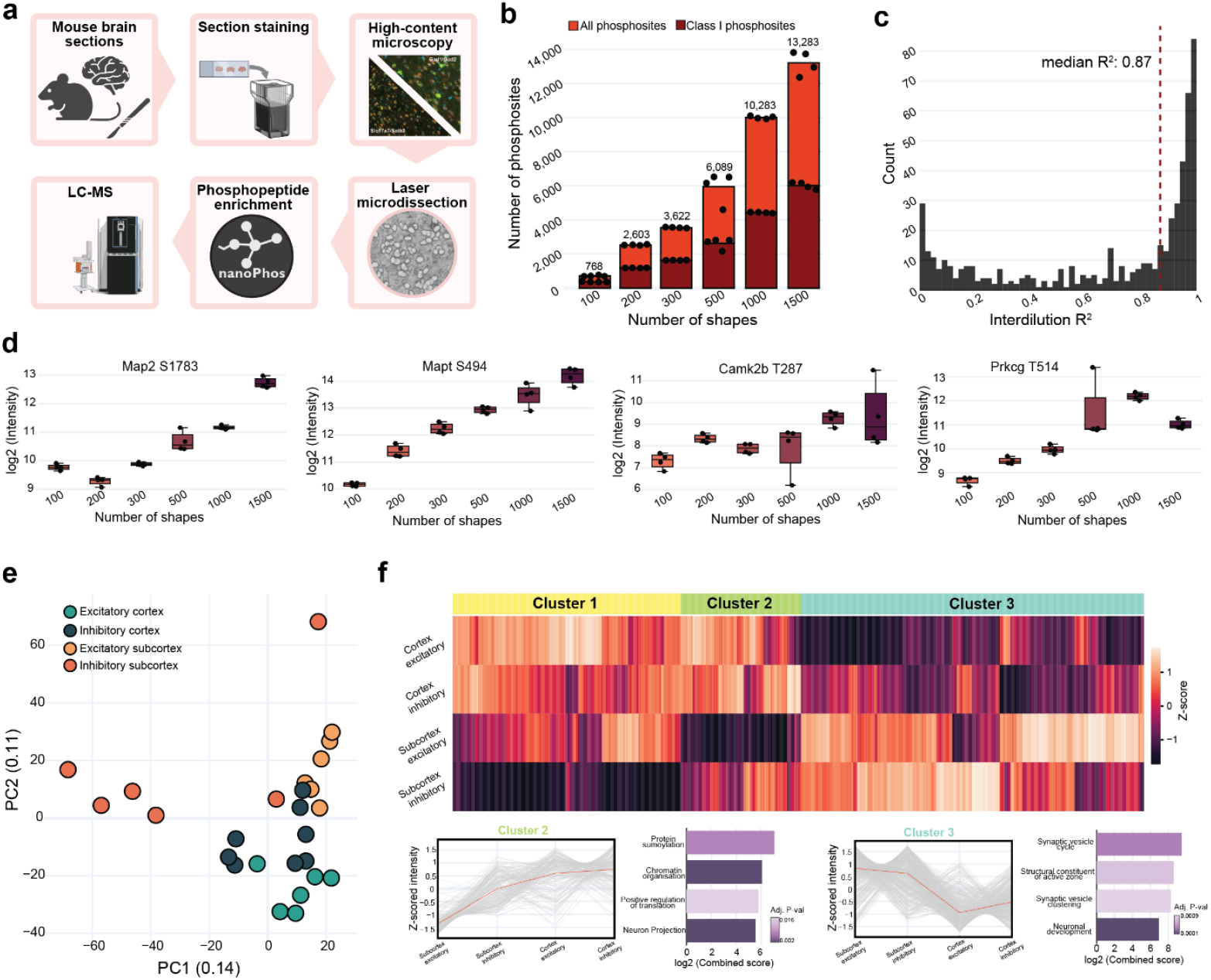
Deep Visual Phosphoproteomics (phosphoDVP) of mouse brain reveals cell type– and region-specific signaling. **a** Schematic overview of Deep Visual Proteomics (DVP) combined with phosphoproteomics analysis (phosphoDVP). **b** Number of identified phosphosites and Class I phosphosites (darker color) as a function of the number of microdissected neuronal shapes (n = 4 per condition). **c** Quantitative linearity across shape input amounts shown as histogram for all phosphosites, identified in at least 3 dilution points. **d** Intensity profiles of selected neuronal phosphosites as a function of the number of excised shapes (n = 4 per condition). **e** PCA of the phosphoproteome for spatially excised neuronal populations (n = 6-8 per condition). **f** Hierarchical clustering heatmap of z-scored intensities for ANOVA-significant phosphosites (FDR < 0.05) across the four neuronal populations. Lower panels show representative intensity profiles and GO enrichment analysis for cluster 2 and cluster 4 (Adj. P-value < 0.05).

We first determined the optimal number of microdissected cellular shapes - the subset of a single cell present in our 10 µm sections - necessary to achieve meaningful phosphoproteomic depth. A dilution series from 100 to 1,500 shapes revealed that phosphosite identifications expectedly scaled with input: from approximately 750 phosphosites with 100 shapes to over 13,000 sites with 1,500 shapes (**Fig. 5b**). Based on protein content estimations from fresh-frozen tissue experiments (**Suppl. Fig. 3a, 3b**), we calculated that 1,000 shapes correspond to approximately 55 ng of protein input - balancing microdissection time and phosphoproteome depth (**Suppl. Fig. 4a**).

Quantitative performance across the dilution series was excellent, with a median interdilution R^2^ of 0.87 (**Fig. 5c**) and robust reproducibility within conditions (**Suppl. Fig. 4b**). Remarkably, even from just 100 shapes - representing approximately 40 neurons - we detected canonical neural phosphorylation sites including Map2 S1783, Mapt S494, Camk2b T287, and Prkcg T514 (**Fig. 5d**). These well-characterized phosphosites showed linear scaling with shape number and remained quantifiable even at the lowest input.

For the main experiment, we collected 1,000 shapes each from four neuronal populations: cortical excitatory neurons, cortical inhibitory neurons, subcortical excitatory neurons, and subcortical inhibitory neurons. To enable normalization of phosphorylation changes against underlying protein abundance, we also collected 100 shapes from each population for proteome analysis. This yielded a comprehensive dataset of 13,000 phosphosites and over 7,600 protein groups (**Suppl. Fig. 4c, d**).

PCA revealed clear separation of neuronal populations by both cell type and anatomical region (**Fig. 5e**). PC1 distinguished excitatory from inhibitory neurons, while PC2 separated cortical from subcortical regions, indicating that both cell identity and anatomical location shape the phosphoproteome. Hierarchical clustering of ANOVA-significant phosphosites revealed four major clusters with distinct regulation patterns (**Fig. 5f**). Cluster 2 showed enrichment in cortical excitatory neurons with pathways related to chromatin organization, protein sumoylation and positive regulation of translation. Cluster 3 was enriched in subcortical regions with terms including synaptic vesicle clustering and neuronal development

Differential comparison between cortical excitatory and inhibitory neurons identified 91 significantly upregulated and 73 downregulated phosphosites in excitatory neurons (**Suppl. Fig. 4e**). Among the most differential sites were phosphorylations on synaptic scaffolding proteins (SHANK1) and ion channels (CACNA1A), reflecting the distinct synaptic architectures of these cell types. These results establish phosphoDVP as a platform for cell type-specific, spatially resolved phosphoproteomics in vivo, linking measurements of molecular signaling to tissue architecture and cellular context.

## Discussion

Here, we describe nanoPhos, a phosphoproteomics workflow that overcomes key limitations of existing methods by leveraging the SPEC protocol for high recovery sample preparation in nanoliter scale volumes. This enables deep, quantitative and robust phosphoproteomics from nanogram-scale protein inputs. Its design prioritizes compatibility with strong detergents, minimal-loss processing, and automation for maximizing phosphosite coverage and enrichment selectivity as well as providing reproducible quantitation. At 10 ng input nanoPhos accurately quantified a number of highly functional phosphorylation sites and achieved nearly hundred-fold higher coverage than the recent µPhos workflow under identical MS conditions. Compared to phosphoproteomics workflows of a decade ago, which generally required 10 mg of input material, nanoPhos on modern LC-MS instrumentation achieves a million-fold sensitivity gain^31^.

The SPEC sample preparation workflow that underlies nanoPhos is described in detail in a companion paper^19^, where we demonstrate its broad applicability to diverse sample types and its compatibility with standard laboratory automation. Together, these workflows establish a modular platform for ultra-sensitive proteomics and phosphoproteomics.

The sensitivity gain provided by nanoPhos has the potential to fundamentally expand the scope of phosphoproteomics research across multiple research areas. First, it enables the deep phosphoproteome profiling from rare cell populations. With as few as 100 FACS-sorted cells yielding nearly 10,000 phosphosites, researchers can now directly interrogate signaling states in rare cell populations that were previously inaccessible to phosphoproteomics. This capability transforms phosphoproteomics from mainly bulk-based technique into a tool suited to dissect cellular heterogeneity. Next, nanoPhos unlocks the vast repositories of archived FFPE clinical samples for phosphoproteomic investigation. Despite the challenges of crosslinking and embedding, we recovered tens of thousands of phosphosites from nanograms of starting protein material with high correlation to fresh-frozen tissue. This opens retrospective studies of signaling pathway alterations across disease progression, treatment response, and patient stratification.

The most impactful advance we present is the integration of nanoPhos with DVP, enabling global phosphoproteomic measurements in spatially and cell-type– resolved tissue contexts. A longstanding goal of spatial biology has been to measure signaling states without mixing cell types, a goal that phosphoDVP now makes possible. This approach bridges molecular signaling and tissue architecture, providing a new technology for understanding disease microenvironments and cellular heterogeneity *in situ*.

The ability to measure phosphorylation events with spatial and cellular resolution opens new possibilities for functional tissue proteomics. In oncology, this could mean directly assessing the signaling state and kinase inhibitor vulnerability of different cancer cell populations, quantifying signaling heterogeneity in immune infiltrates or cancer-associated stromal cells, or identifying drug-resistant clones within otherwise responsive lesions. In neuroscience, phosphoDVP could help decode synapse-specific phosphorylation dynamics or capture drug-induced signaling responses in brain cell types and regions^8^. In developmental biology, it could reveal how morphogen gradients translate into cell-type-specific phosphorylation patterns during tissue patterning. Combined with multiplexed imaging and AI-driven phenotyping, spatial proteomics will no longer be limited to protein levels, but will include the dynamic regulation that defines cell state and fate.

Several limitations should be noted. First, nanoPhos currently requires at least 100 cells or about 10 ng of input for robust phosphoproteome coverage; true single-cell phosphoproteomics remains out of reach. Second, the efficiency of enrichment of low-stoichiometry modifications - particularly phosphotyrosine - still remains less effective as the enrichment of phosphoserines and phospho-threonines and requires dedicated protocols at low-microgram and sub-microgram inputs. Third, as with all LMD-based approaches, phosphoDVP throughput is constrained by microdissection time, making it best suited for targeted studies of defined cell populations rather than unbiased tissue-wide surveys. Future developments in automated microdissection, single-cell sample preparation, and MS sensitivity may address these constraints.

In summary, nanoPhos provides a robust, sensitive, and automatable platform for phosphoproteomics at the nanogram scale. Its integration with DVP extends spatial proteomics to post-translational modifications, enabling cell-type-resolved signaling analysis in intact tissue. We anticipate that phosphoDVP will find broad application in neuroscience, oncology, and developmental biology, wherever understanding signaling in spatial and cellular context is essential.

## Acknowledgements

We thank our colleagues at the Department of Proteomics and Signal Transduction at the Max Planck Institute of Biochemistry. In particular, we would like to thank Bianca Splettstößer and Igor Paron for technical support as well as Medini Steger for administrative support. FACS support was provided by Martin Spitaler and Markus Oster at the Imaging Core Facility at the Max Planck Institute of Biochemistry.

## Code availability

All code that was used to analyze data and generate figures can be found at https://github.com/DenysOliinyk3007/nanoPhos-figures

## Ethical approval

The study was conducted in accordance with the ethical principles for medical research of the Declaration of Helsinki and was approved by the Ethics Committee of the Charité University Medical Department in Berlin (EA4/082/22).

## Methods

### Human cell culture

Human epithelial carcinoma cells of the line HeLa (ATCC, S3 subclone) were cultured in Dulbecco’s modified Eagle’s medium containing 20 mM glutamine, 10% fetal bovine serum, and 1% penicillin-streptomycin. Cells were routinely tested for mycoplasma contamination. For dilution series experiments, HeLa cells were cultured until 80% confluency, harvested with 0.25% trypsin/EDTA and collected in 15 mL falcon tubes. Cells were then washed twice with cold TBS and pelleted by centrifugation at 200*g* for 10 min. Next, supernatant was aspirated, cells were snap-frozen in liquid nitrogen and stored until further use. For EGF experiments, Hela cells at a plate confluence of 80% were serum-starved for 3 hours and subsequently treated for 15 min with 125 ng/mL animal-free recombinant human EGF or distilled water. After treatment, cells were washed three times with ice-cold TBS, snap-frozen in liquid nitrogen and stored in −80 °C until further use.

### Mouse embryonic stem cell culture

WT ground state E14 mESCs were cultured in serum-free medium consisting of: N2B27 (50% neurobasal medium (Life Technologies), 50% DMEM/F12 (Life Technologies)), 2i (1 μM PD032591 and 3 μM CHIR99021 (Axon Medchem, Netherlands)), 1000 U/ml recombinant leukemia inhibitory factor (LIF, Millipore), and 0.3% BSA (Gibco), 2 mM L-glutamine (Life Technologies), 0.1 mM β-mercaptoethanol (Life Technologies), N2 supplement (Life Technologies), B27 serum-free supplement (Life Technologies), and 100 U/ml penicillin, 100 μg/ml streptomycin (Sigma). Ground state mESCs were transferred to serum + LIF conditions (S+L) consisting of Dulbecco’s modified Eagle’s medium (DMEM, Sigma) supplemented with 16% fetal bovine serum (FBS, Sigma), 0.1 mM ß-mercaptoethanol (Life Technologies), 2 mM L-glutamine (Sigma), 1× MEM Non-essential amino acids (Sigma), 100 U/ml penicillin, 100 μg/ml streptomycin (Sigma) and 1000 U/ml recombinant leukemia inhibitory factor (LIF, Millipore) and adapted to this medium for at least 7 days. For the retinoic acid differentiation, S+L mESCs were transferred to RA medium consisting of DMEM supplemented with 16% FBS, 0.1 mM ß-mercaptoethanol (Life Technologies), 2 mM L-glutamine (Sigma), 1× MEM Non-essential amino acids (Sigma), 100 U/ml penicillin, 100 μg/ml streptomycin (Sigma) and 1 µM retinoic acid (bio techne) for 48 hours.

### FACS of HeLa and mESC cultured cells

Cells were cultured until 80% confluency, counted and harvested with 0.25% trypsin/EDTA (for HeLa cells) to 15 mL falcon tubes. Cells were then washed twice with cold TBS, pelleted by centrifugation at 200*g* for 10 min and resuspended in TBS to achieve concentration of 1 million cells per 1 mL. Subsequently, 1 µL of DAPI was added to cell suspension and FACS was performed on DAPI-negative live cell population in the ‘Purity’ sorting mode. Cells were sorted into 384-well TwinTec Eppendorf plates containing 7 µL of lysis buffer (2% SDC, 0.1% DDM, 10 mM TCEP, 40 mM CAA in 100 mM Tris-HCl, pH 8.5), sealed with aluminum foil, centrifuged briefly and frozen at −80 °C until further use.

### Mouse experiments

Eight-week-old female mice of genetic background C57BL/6J were used for excitatory and inhibitory neuron analysis. Animals used were bred for scientific purposes, and the research in this project does not involve experiments on animals (as defined by law). All animals were sacrificed by CO_2_ euthanasia prior to removal of brains in accordance with the European Commission Recommendations for the euthanasia of experimental animals (Part 1 and Part 2). Breeding, housing, and euthanasia of the animals are fully compliant with all German (i.e., German Animal Welfare Act) and EU (i.e., Directive 2010/63/EU) applicable laws and regulations concerning care and use of laboratory animals.

### Organ collection and immunofluorescence staining for phosphoDVP experiments

After euthanasia, brains were dissected and embedded in Neg-50 (epredia). 10um coronal cryosections were collected onto 2um PEN membrane slides (MicroDissect GmbH). Slides were stained with NucGreen (Thermo) 1:100 in PBS and HCR v3.0 probes to Slc17a7 and Satb2 (excitatory neurons) or Gad1 and Gad2 (inhibitory neurons) following the standard HCR v3.0 protocol (PMID: 29945988). HCR fluorescent amplifiers B2-546 and B4-647 were used for Slc17a7 and Satb2 probes, respectively, or for Gad1 and Gad2 probes, respectively.

### High-content imaging and image processing for phosphoDVP experiments

Imaging was performed on the Axioscan 7 slide scanner (Zeiss) equipped with Colibri 7 LED light source and appropriate filter sets (for 488, 546, and 647nm channels). A 20x NA 0.8 Plan-Apochromat objective was used. Z stacks were processed to single Z-planes with software Zen 3.7 (Zeiss) using the Extended Depth of Focus variance method, and then image tiles were stitched using the Zen stitching function. Stitched images were imported into Biological Image Analysis Software (BIAS, Single-Cell Technologies), and segmentation was carried out on the nuclear channel with Cellpose v2.3.2 and masks imported into BIAS. Brain images were hand-annotated for cortical vs sub-cortical regions, and double-positive cells (either for excitatory or inhibitory markers) from cortical vs sub-cortical regions were selected for laser microdissection.

### Laser microdissection for phosphoDVP experiments

Contour coordinates were imported, and shapes cut using the LMD7 (Leica) laser microdissection system in a semi-automated mode with the following settings: power 55; aperture 1; speed 75; middle pulse count 1; final pulse 0; head current 45 – 50%; pulse frequency 2.9 and offset 190. The microscope was operated with the LMD v8.5.9136 software, and samples collected into 384-well plates, leaving the outmost rows and columns empty. Plates were then sealed, centrifuged at 3,000*g* for 3 min, and frozen at −20 °C for further processing.

### Slide processing for microbulk phosphoproteomics

We assembled a cohort of 10 lung adenocarcinoma cases (ethics approval # EA4/082/22). We pretreated slides with VECTABOND as follows: slides were exposed to UV light (254 nm) for 1 hour, immersed in 100% acetone for 5 minutes, and coated with VECTABOND solution (VECTABOND Reagent, Vector Laboratories; 7 mL reagent in 350 mL acetone) for 5 minutes. We then washed slides three times by dipping in distilled water and allowed them to dry. 10 μm sections were cut from formalin-fixed paraffin-embedded (FFPE) tissue blocks using a microtome (Microm HM 340E, Thermo Fisher Scientific) and mounted onto the pretreated 1.0 PEN membrane slides (MicroDissect). We dried mounted sections overnight at 37°C. We then heated slides at 55°C for 30 minutes, deparaffinized them in xylene (2 × 2 minutes), followed by rehydration through a graded ethanol series: 100% ethanol (2 × 1 minute), 95% ethanol (2 × 1 minute), 75% ethanol (2 × 1 minute), 30% ethanol (2 × 1 minute), and double-distilled water (ddH_2_O; 2 × 1 minute). For antigen retrieval and decrosslinking, we incubated slides in a buffer containing 10% glycerol (v/v) in Dako Target Retrieval Solution pH 9 (Agilent) at 90°C for 30 minutes in a water bath. We allowed slides to cool for 15 minutes at room temperature, washed them twice in ddH_2_O (5 minutes each), and airdried them before laser microdissection.

### Laser microdissection for microbulk experiments

We performed laser microdissection using an LMD7 microscope (Leica Microsystems) with a 10× objective. We collected tissue areas into 384-well plates (Eppendorf twin.tec PCR Plate 384 LoBind). For phosphoproteomics analysis, we collected 200,000 μm^2^ of tissue material per sample; for total proteomics analysis, we collected 100,000 μm^2^ per sample according to pathologist annotations. For each sample, we excised multiple circular areas (25,000 μm^2^ each) from regions of interest until reaching the target area (8 areas for phosphoproteomics, 4 areas for total proteomics). For each case, we selected annotated adenocarcinoma and adjacent healthy lung tissue areas and cut 6 samples per area (3 for phospho- and 3 for proteome analysis) per case for total of 120 samples (**Supplementary Table 3**). The following laser parameters were used: power = 55, aperture = 13, speed = 4, middle pulse count = 4, final pulse = 25, head current = 74%, pulse frequency = 3446 Hz, offset = 50, operating in middle pulse cutting mode. After collection, 384- well plates were centrifuges at 2,000*g* for 10 minutes and stored at room temperature until further processing or directly transferred onto preequilibrated SPEC tips.

### Cell lysis for cell culture experiments

For bulk cell lysate experiments, frozen HeLa cell pellets were resuspended in a lysis buffer (2% SDC, 0.1% DDM, 10 mM TCEP, 40 mM CAA in 100 mM Tris-HCl, pH 8.5) and boiled for 15 min at 95 °C while mixing at 1500 rpm. This was followed by high-energy tip sonication (10 pulses, 5 sec on, 5 sec off, 20% duty cycle). Lysates were then centrifuged for 5 min at maximum speed to remove cell debris. Protein concentration was determined via tryptophan assay. Samples were then diluted with 0.5% SDC in 100 mM Tris-HCl to the input concentration of 1 µg/µl and frozen at −80 °C until further processing or directly transferred onto preequilibrated SPEC tips.

### Cell lysis for FACS-related experiments

Cell-containing 384-well plates were incubated for 15 min at 95 °C in a 384-well thermal cycler (Eppendorf) at a lid temperature of 110 °C. This was followed by brief centrifugation and sonication for 10 minutes in a sonication water bath. Plates were then briefly centrifuged and subjected to another 15 min of incubation at 95 °C in a 384-well thermal cycler. Finally, plates were centrifuged for 2 min at 2,000*g* and stored at −80 °C until further processing or directly transferred onto preequilibrated SPEC tips.

### Tissue lysis for bulk dilution experiments

FFPE tissue samples were deparaffinized by incubating approximately 300 µg of sample in 300 µL n-Heptane for 1min at 30 °C and 700 rpm, discarding the solvent, repeating this step once more with n-Heptane, and two more times with 300 µl methanol. Deparaffinized FFPE and fresh-frozen tissues were resuspended in a lysis buffer (2% SDC, 0.1% DDM, 10 mM TCEP, 40 mM CAA in 100 mM Tris-HCl, pH 8.5) and boiled for 5 min at 95 °C, followed by tip-sonication (10 pulses, 5 sec on, 5 sec off, 20% duty cycle). The samples were then boiled again for 5 min at 95 °C. Protein concentration was then determined using the tryptophan assay. The sample was then diluted with 0.5% SDC in 100 mM Tris-HCl to the input concentration of 1 µg/µl and stored at −80 °C until further processing or directly transferred onto preequilibrated SPEC tips.

### Sample preparation of phosphoDVP and microbulk phosphoproteomics samples

All liquid handling steps were performed on a Bravo pipetting robot as described before^15^. During each incubation step, plates were tightly sealed with two layers of sealing aluminum foil to avoid evaporation. Shape-containing 384-well TwinTec Eppendorf plates were retrieved from the - 80 °C and centrifuged at 3,000*g* for 2 min. The wells were then washed on the robot with 28 µl of 100% ACN and dried in a SpeedVac (Eppendorf) at 30 °C for 45 min. Shapes were then resuspended in 7 µl of lysis buffer (2% SDC, 0.1% DDM, 10 mM TCEP, 40 mM CAA in 100 mM Tris-HCl, pH 8.5) and incubated for 15 min at 95 °C in a 384-well thermal cycler (Eppendorf) at a lid temperature of 110 °C. This was followed by brief centrifugation and sonication for 10 minutes in a sonication water bath. Plates were then briefly centrifuged and subjected to another 15 min of incubation at 95 °C in a 384-well thermal cycler. Finally, plates were centrifuged for 2 min at 2,000*g* and protein lysates were transferred to preequilibrated SPEC tips.

### Sample preparation of DVP and microbulk samples for proteomics analysis

All liquid handling steps were performed on a Bravo pipetting robot. Samples were collected into 384-well TwinTec Eppendorf plates and prepared via our standard DVP workflow. Briefly, samples were lysed in 7 µl of 70mM TEAB and 0.013% DDM for 60 min at 95 °C. Next, 1 µL of 100% ACN was added to each well and plate next boiled for additional 60 min at 72 °C. Proteins were proteolyzed overnight with LysC and Trypsin at 37 °C in a 384-well thermal cycler (Eppendorf) at a lid temperature 50 °C. Resulting peptides were then acidified with 10% TFA and loaded onto preequilibrated EvoTips. Microbulk proteome samples were prepared using a SPEC workflow (see below).

### SPEC workflow

SPEC tips were prepared by placing two plugs of strong-anion-exchange (SAX) material (3M Empore, catalog # 13-110- 024) in a pipette tip with a blunt-ended syringe needle. Before sample loading, SPEC tips were activated with 10 µl 100% dimethylsulfoxide (DMSO) and centrifugation at 700*g* for 1.5 min. Next, tips were preequilibrated with 20 µl SPEC Loading buffer (20 mM CAPS, 0.01% DDM in ddH_2_O) and centrifuged at 700*g* for 2 min. Protein sample was then alkalinized by adding to SPEC Loading buffer in a ratio 1:10 and loaded on SAX material by centrifugation at 500*g* for 10 min. Tips were then washed with 20 µl SPEC Washing buffer (50 mM TEAB, 0.01% DDM in ddH_2_O) and centrifuged at 700*g* for 2 min or until liquid is fully through. Finally, proteins were on-tip digested by adding 2 µl of digestion buffer (25 ng/µl trypsin/LysC mix in 50 mM TEAB) and centrifuging for 20 sec at 100*g*. Tips were then incubated at 37 °C for 60 min. For proteome measurements peptides were eluted with 20 µl of SPEC Elution buffer (1% FA, 0.01% DDM in ddH_2_O) and centrifugation at 300*g* for 5 min onto preequilibrated Evotips. For phosphoproteome analysis peptides were eluted by the addition of 20 µl of nanoPhos Elution buffer (1M NaCl, 10% ACN, 0.1% DDM, 1% FA in ddH_2_O) to a 96-well TwinTec Eppendorf plate, which already contained 80 µl of nanoPhos preelution buffer (90% ACN, 0.1% DDM, 1% FA in ddH_2_O).

### Phosphopeptide enrichment for nanoPhos experiments

Phosphopeptide enrichment cartridges, each containing 5 µl Fe(III)-nitrilotriacetic acid (Agilent, part number: G5496-60085), were first primed with 100 µl priming buffer (1% FA, 99% ACN) with a flow rate of 300 µl/min, followed by equilibration with 50 µl wash/equilibration buffer (1% FA, 80% ACN in ddH_2_O) with a flow rate of 5 µl/min. Peptides were then loaded on cartridges with a flow rate of 5 µl/min and subsequently washed with 50 µl wash/equilibration buffer with a flow rate of 5 µl/min. Phosphopeptides were eluted with 25 µl of elution buffer (500 mM NH_4_H_2_PO_4_ in ddH_2_O) with a flow rate of 5 µl/min directly onto preequilibrated EvoTips.

### µPhos workflow

µPhos phosphopeptide enrichment was performed as described before^10^. Briefly, HeLa protein lysates were transferred to 96- well deep-well plates (Eppendorf) and diluted with the lysis buffer to 19 µl. 1 µl of digestion buffer was added to each well. The plate was then sealed with a silicone mat and incubated for 2 hours at 1,500 rpm at 37 °C. After digestion, plate was briefly centrifuged and 20 µl of 100% 2-propanol was added and plate was incubated for 30 sec at 1,500 rpm at room temperature, followed by addition of 40 µl of µPhos Enrichment Buffer. Next, 5 µl of 1 mg/µl TiO_2_ solution was added to peptides, after which plate was incubated at 40 °C at 1,500 rpm for 7 min. Plate was then centrifuges and supernatant was aspirated with a multi- channel pipette. Beads were then washed five times with 200 µl of µPhos Washing Buffer. Next, beads were transferred to C8 StageTips and centrifuged at 700*g* for 7 min. Phosphopeptides were then eluted by two-step addition of 30 µl of µPhos Elution Buffer and centrifugation for 4 min at 700*g*. Eluates were then vacuum dried for 30 min at 45 °C until <10 µl was left. 200 µl of Evosep buffer A (0.1% FA in ddH_2_O) was then added to eluates and solution was transferred on preequilibrated EvoTips.

### Peptide loading of C-18 tips

C-18 tips (Evotip Pure, Evosep) were washed once with 20 µl of buffer B (99.9% ACN, 0.1% FA), activated for 1 min in 2- propanol and equilibrated with 20 µl of buffer A. Phosphopeptides were eluted into 225 µl of buffer A in the tip, which was then centrifuged for a few seconds. After peptide binding, the disk was further washed once with 20 µl buffer A and further overlayed with 150 µl buffer A. All centrifugation steps were performed at 700*g* for 1 min, except sample loading for 3 min.

### LC-MS/MS analysis

The samples were analyzed using an Evosep One LC system (Evosep) coupled to an Orbitrap Astral Zoom mass spectrometer (Thermo Fisher Scientific). Peptides were eluted from the Evotips using a ‘Whisper Zoom’ gradient with a throughput of 80 samples per day on an Aurora Rapid column of 5-cm length, 75- µm-internal diameter, packed with 1.7 µm C18 beads (IonOpticks). The column temperature was maintained at 60 °C using a column heater (IonOpticks). The Orbitrap Astral Zoom was equipped with an EASY-Spray source (Thermo Fisher Scientific). An electrospray voltage of 1,900 V was applied for ionization, and the radio frequency level was set to 40. For phosphoproteomics samples Orbitrap MS1 spectra were acquired from 380 to 1,380 m/z at a resolution of 240,000 (at m/z 200) with a normalized automated gain control (AGC) target at 500% and a maximum injection time of 3 ms. For the Astral MS/MS scans in data-independent acquisition (DIA) mode, we used 100 variable isolation windows, designed with a pyDIAid software^21^ (**Supplementary Table 1**). A maximum injection time of 10 ms was used. The isolated ions were fragmented using high-energy collisional dissociation with 25% normalized collision energy. For proteomics samples Orbitrap MS1 spectra were acquired from 380 to 980 m/z at a resolution of 240,000 (at m/z 200) with a normalized automated gain control (AGC) target at 500% and a maximum injection time of 3 ms. For the Astral MS/MS scans in data-independent acquisition (DIA) mode, 200 equidistant isolation windows of 3 Th each were used with maximum injection time of 6 ms (**Supplementary Table 2**). The isolated ions were fragmented using high-energy collisional dissociation with 25% normalized collision energy. Samples were acquired in randomized order within each condition.

### Spectral search

LC-MS raw files were processed in Spectronaut v20.1 without experimental spectrum libraries (‘directDIA+’ workflow in Spectronaut). Data were searched against the UniProt human or mouse reference proteome (accessed August 2024). We set the protease specificity to trypsin with a maximum number of two missed cleavages and required a minimum peptide length of 7 amino acids. The mass tolerances for precursor and fragment ions were set to ‘Dynamic’ for both MS1 and MS2 level. False discovery rates were controlled by a target-decoy approach to ≤1% at precursor and protein levels. For phosphoproteomics experiments, we defined cysteine carbamidomethylation as a fixed modification and protein N-terminal acetylation, methionine oxidation and serine/threonine/tyrosine (STY) phosphorylation as variable modifications in ‘BGS Phospho PTM Workflow’ and activated the PTM localization mode. For proteomics runs we used ‘BGS Factory Settings (default)’ workflow with default settings. To report all identified phosphopeptides, we defined a localization probability score threshold of 0 and, if applicable, filtered the output on the phosphosite level as described below. Quantification values were filtered by q-value and we defined the ‘Automatic’ normalization mode for cross-run normalization.

### Bioinformatics data analysis

Bioinformatics data analysis was carried out in the Python programming environment (v. 3.11). For subsequent statistical analysis, phosphoproteomics raw tabular data was exported from Spectronaut using ‘BGS Factory Report’ export schema with ‘EG.PrecursorID’, ‘PEP.PeptidePosition’, ‘EG.PTMAssayProbability’, ‘PG.Genes’ and ‘PG.ProteinGroups’ as additional columns. This was next parsed with a custom Python implementation of the ‘PeptideCollapse’ plugin for Perseus^32^. Phosphopeptide enrichment selectivity was parsed from Spectronaut software as the percentage of all identified precursors that are carrying a phosphorylation modification. Proteome datasets were exported from Spectronaut using ‘Protein Quant Pivot Report’ export schema. Protein or phosphosite intensities were log_2_- transformed and average intensities of the biological or technical replicates were computed for each analyzed condition where applicable. For differential analysis, phosphosites with localization probability < 0.75 and those phosphosites which exhibited an identification rate of < 70% were removed. Furthermore, where applicable, samples with identification rates less than 1.5x the interquartile range from the median experiment identification rate were removed. Imputation of the remaining missing values was performed based on a normal distribution (width of 0.3, downshift of 1.8). PCA analysis was performed on an imputed dataset with ‘scikit-learn’ python library (version 1.6.1). Where applicable, phosphoproteomics datasets were normalized by subtracting from each phosphosite the median abundance of its corresponding parent protein within each condition. Box plots show the median (center line) with interquartile range of 25% to 75%, whiskers extend to furthest data points within 1.5x the interquartile range from the box boundaries. For the ANOVA, the ‘pingouin’ python package (version 0.5.5) was used and intensities were z-scored prior to analysis. Subsequent multiple testing correction was conducted using Benjamini-Hochberg method with FDR cut-off of 5%. For differential expression, t-test analysis was performed either with ‘pingouin’ python package or with ‘AlphaQuant’ python library (version 0.2.0). For both methods, multiple testing correction was performed (Benjamini-Hochberg method, FDR cutoff 5%). Unsupervised hierarchical clustering was performed on ANOVA-significant proteome-normalized phosphosites and gene ontology analysis was performed with ‘gseapy’ python library (version 1.1.10) and ‘EnrichR’ software (Adj. P-value < 0.05 for both)^33,34^. Kinase enrichment analysis was performed using ‘kinase-library’ python package (version 1.5.0) on phosphosites that were considered statistically up- or downregulated by t-test analysis (Adj. P-value < 0.05, |log2FC| > 0.585)^35,36^. Interdilution R^2^ was determined by fitting a linear regression to each phosphosite that was identified in at least 3 points of a dilution curve and calculating a coefficient of determination (R^2^). Mouse signaling pathways for **Fig. 4e** were derived by annotating identified phosphorylated proteins with an annotation database, containing GOBP, GOCC, GOMF and KEGG terms. For estimation of protein input amounts for phosphoDVP analysis, we fitted a logarithmic regression curve for bulk mouse fresh-frozen tissue and linear regression curve for bulk mouse FFPE tissue and derived fit equations (**Suppl. Fig. 3a, 3b**). All figures were plotted with ‘plotly’ python library. Icons for **Fig. 1, Fig. 2a** and **Fig. 4d** were retrieved from Biorender and manually adjusted in Adobe Illustrator.

## Supplementary Materials

**Supplementary Figure 1.**
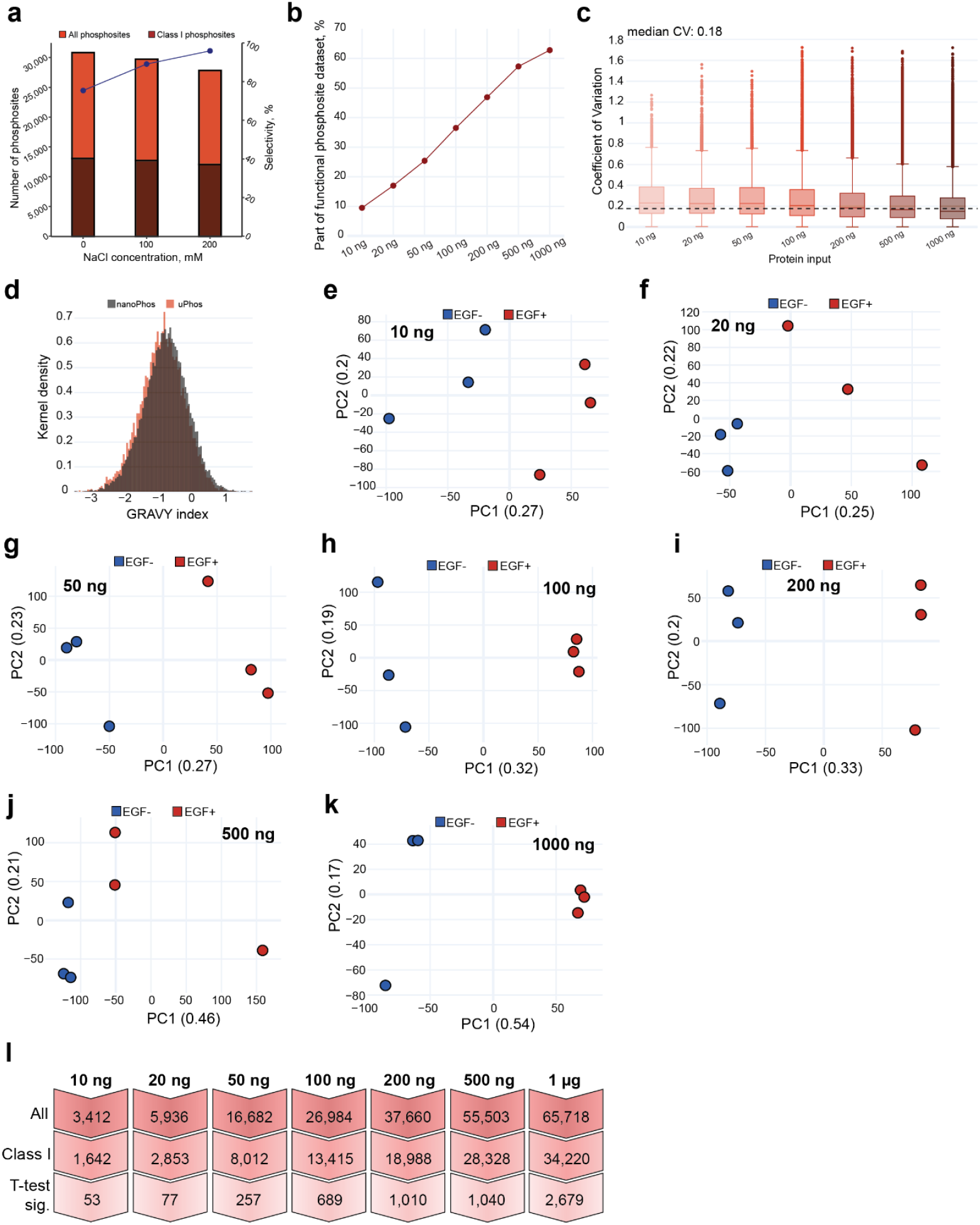
**a** Number of unique phosphorylation sites and phosphopeptide enrichment selectivity as a function of NaCl concentration in phospho-enrichment buffer (n = 1) **b** Percentage of functional phosphosites (functional score > 0.5 from Ochoa et al., 2020) identified as a function of HeLa protein input amount across the HeLa dilution series. **c** Precision of label-free phosphopeptide quantification shown as coefficient of variation across workflow replicates for the HeLa lysate dilution series (n = 3 per condition). **d** Overlay of GRAVY hydrophobicity index distributions for phosphopeptides detected by nanoPhos (black) versus those detected by µPhos (red). **g** Quantitative reproducibility across the EGF dilution series shown as histogram of interdilution R^2^ values for all phosphosites identified in at least 3 conditions. **e-k** PCA of EGF-treated (EGF+, red) and untreated (EGF-, blue) HeLa samples at different protein input amounts: **e** 10 ng, **f** 20 ng, **g** 50 ng, **h** 100 ng, **i** 200 ng, **j** 500 ng, and **k** 1000 ng (1 µg) (n = 3 per condition). **l** Number of all phosphosites, Class I phosphosites, and t-test significant phosphosites (Adj. P-value < 0.05, |log2FC| > 1) identified across the EGF stimulation dilution series from 10 ng to 1 µg HeLa lysate input series.

**Supplementary Figure 2.**
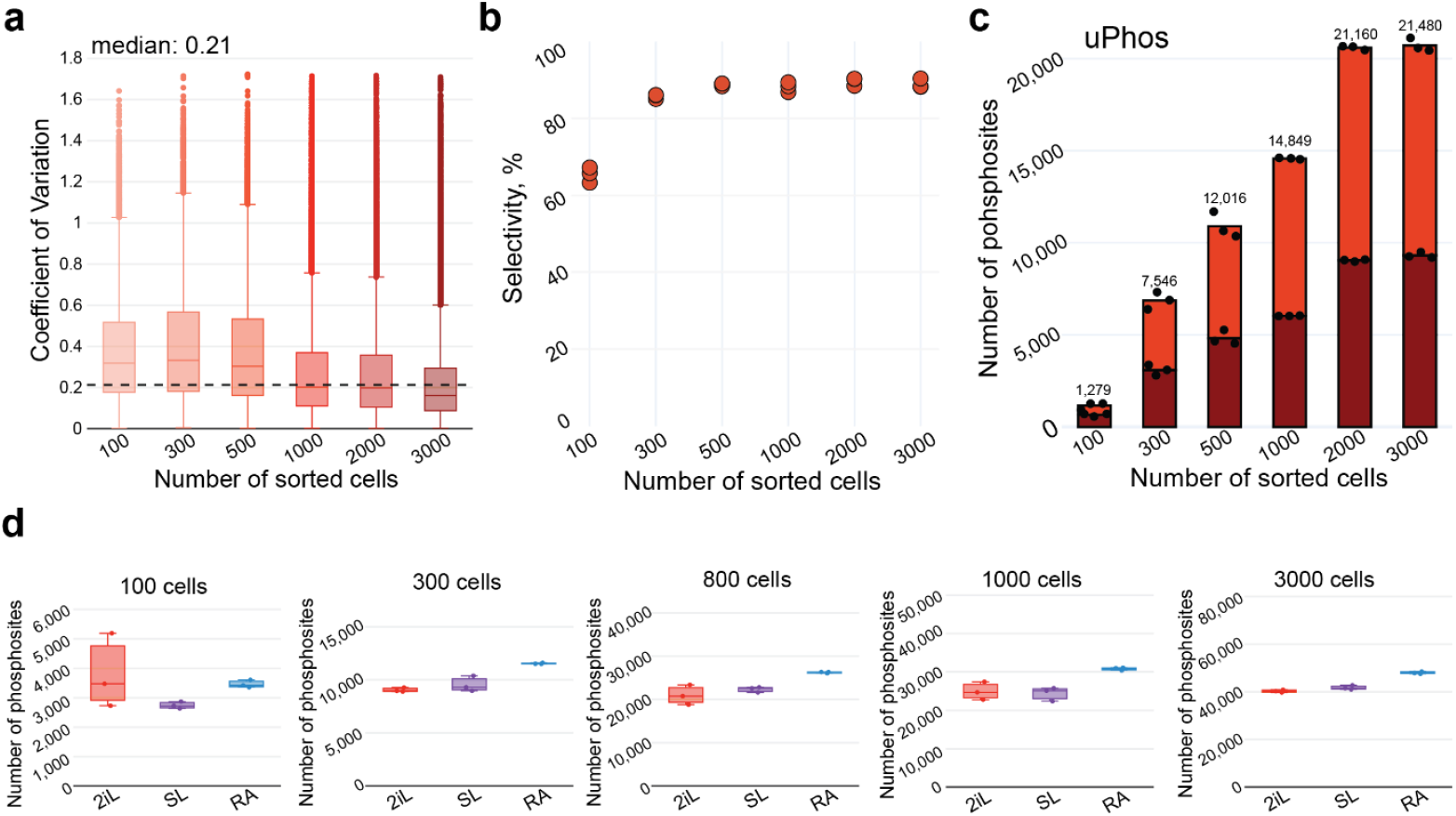
**a** Coefficient of variation across technical replicates (*n* = 3 per condition) for the FACS-sorted cell series, processed with nanoPhos. **b** Selectivity of phosphopeptide enrichment in percent across the sorted cell dilution series (n = 3 per condition). **c** Number of identified phosphosites and Class I as a function of FACS-sorted HeLa cell number identified with uPhos workflow (n = 3 per condition). **d** Number of identified phosphosites from three mESC conditions (2iL = red, SL = purple, RA = blue) across the sorted cell dilution series from 100 to 3,000 cells, shown as box plots for each input amount (n = 3 per condition).

**Supplementary Figure 3.**
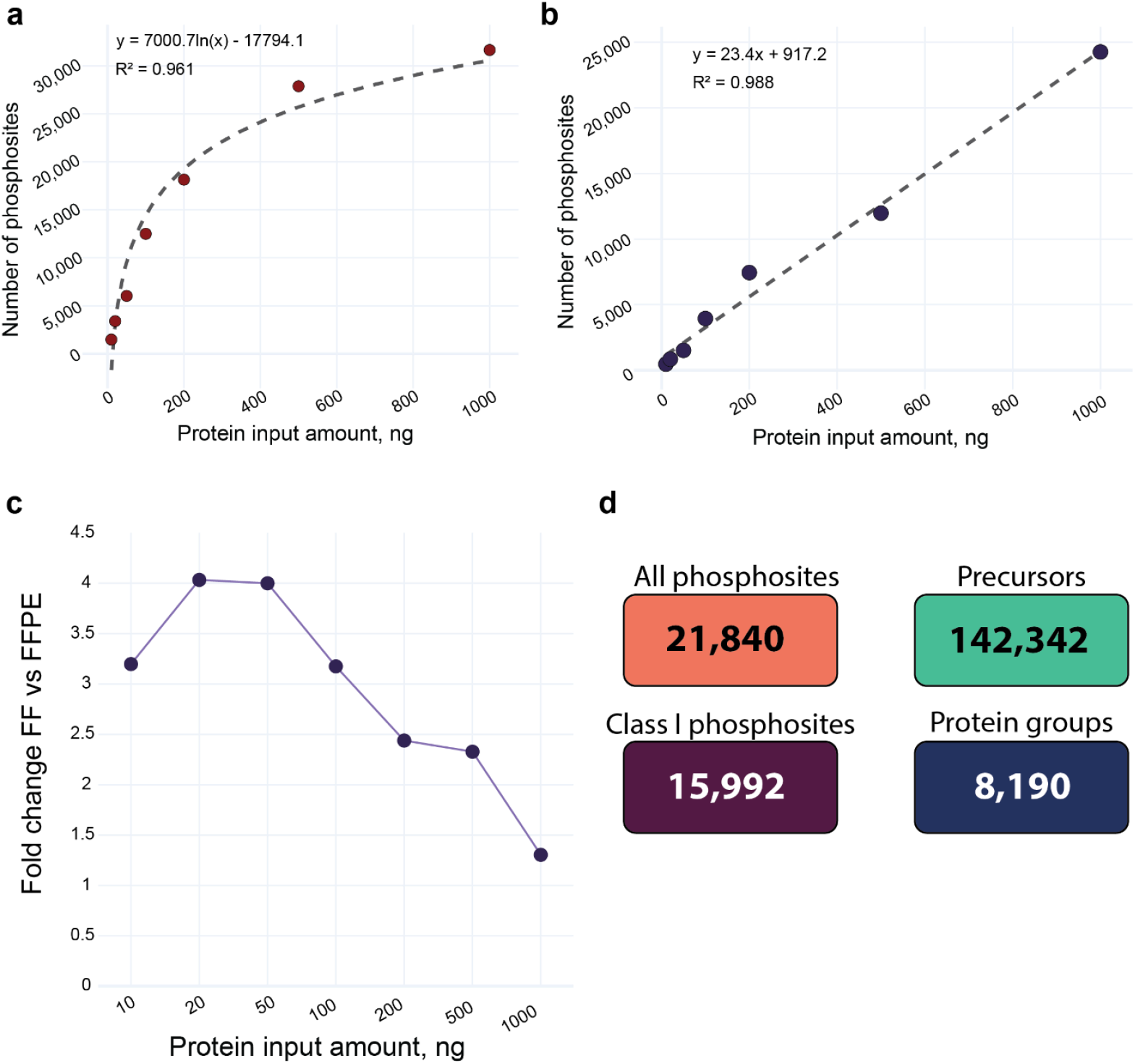
**a** Regression analysis of phosphosite identifications as a function of protein input amount for fresh-frozen mouse brain tissue. **b** Same as **a** but for FFPE mouse brain tissue. **c** Fold change difference in mean phosphosite identifications between fresh-frozen and FFPE tissue as a function of protein input amount. **d** Number of phosphosites, class I phosphosites, peptide precursors and protein groups identified in microbulk adenocarcinoma experiment.

**Supplementary Figure 4.**
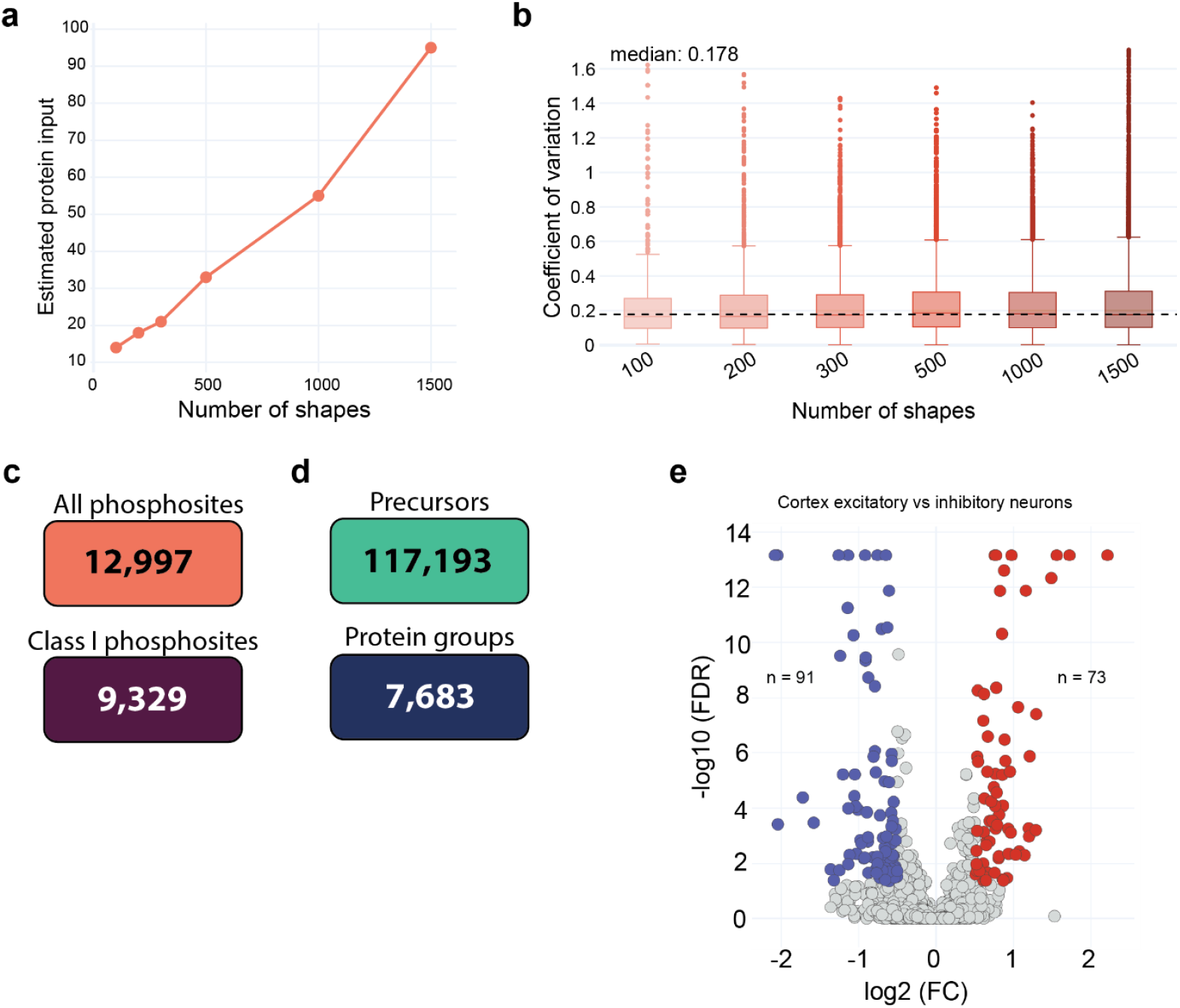
**a** Estimated protein input per number of shapes based on regression analysis from fresh-frozen tissue experiments. **b** Precision of phosphopeptide quantification across the shape dilution series shown as coefficient of variation for replicates (n = 3 per condition). **c** Number of phosphosites and class I phosphosites identified in the main phosphoDVP experiment. **d** Same as **c** but for the number of peptide precursors and protein groups. **e** Volcano plot (FDR < 0.05, FC > 1) of differential phosphosites regulation between cortical excitatory and inhibitory neurons.

